# Gfa2bin enables graph-based GWAS by converting genome graphs to pan-genomic genotypes

**DOI:** 10.1101/2024.12.05.626966

**Authors:** Sebastian Vorbrugg, Ilja Bezrukov, Zhigui Bao, Wenfei Xian, Detlef Weigel

## Abstract

Variation graphs offer superior representation of genomic diversity compared to traditional linear reference genomes, capturing complex features that are otherwise inaccessible to analysis. It seems self-evident that integrating these graphs with genome-wide association studies (GWAS) should enable more comprehensive understanding of genetic landscapes, potentially uncovering novel associations between genetic variations and traits. This approach takes full advantage of rich genomic information, thereby providing deeper insights into the genetic base of complex traits. Our tool, *gfa2bin*, offers multiple methods to (i) genotype variation graphs and (ii) convert the genotypes to well-established data formats for genome-wide association studies (GWAS). We demonstrate that variation graphs are feasible alternatives to traditional linear references for GWAS. Our case study using *Arabidopsis thaliana* and 1,695 traits shows that our approach complements SNP-based approaches, often identifying additional associations, with all associations having on average higher significance compared to SNP-based approaches. *gfa2bin* is implemented in Rust. Commented source code is available under MIT license at https://github.com/MoinSebi/gfa2bin. Examples of how to run *gfa2bin* are provided in the documentation. We added several Python scripts and a Snakemake pipeline for easy processing of our tool using larger data sets. In addition, we recommend using packing (https://github.com/MoinSebi/packing) for reduced storage and preprocessing (normalization) of sequence-to-graph alignments coverage.

## Introduction

The advent of long-read sequencing technologies has revolutionized the field of genomics, enabling the generation of more comprehensive and accurate genome information from multiple members of a species at a reasonable cost. These long-read datasets can be assembled into complete genomes, enabling more accurate determination of genome differences than was possible with short-read data(Koren et al. 2018; Luo, Kang, and Schönhuth 2021; Cheng et al. 2021). The transition to long-read sequencing and complete genomes is overcoming the limitations of fragmented and incomplete representations, and is beginning to provide a more comprehensive view of genomic variation, including better haplotype information. Variation graphs have emerged as a promising alternative to traditional linear reference genomes, which often fail to capture the full spectrum of genetic variation present within and across populations or species (Liao et al. 2023).

By encoding non-reference alleles that include structural variants and complex genomic features, variation graphs offer a less biased representation of genetic diversity compared to a collection of linear genomes that have been aligned to a single reference genome (Garrison et al. 2018, 2023; Hickey et al. 2023). This graph-based representation facilitates a wide range of analyses, such as genotyping, variant calling, and population genetics studies across diverse organisms (Sirén et al. 2020; Hickey et al. 2020; Liu et al. 2020).

The emergence of variation graphs presents an exciting opportunity to more fully exploit genomic data. As stated above, graphs offer a more comprehensive depiction of genetic diversity by themselves, but can also be used as a variation-aware reference for aligning short-or long-read data sets. New approaches have been implemented for sequence-to-graph alignments, increasing mapping accuracy and sensitivity in complex regions (Garrison et al. 2018). Such an advanced data structure, either directly or with additional alignment, enables principled genome-wide association studies (GWAS) that go considerably beyond single nucleotide polymorphisms (SNPs) or variants linked to SNPs, thereby furthering the understanding of the entire spectrum of genetic differences responsible for phenotypic variation in a population.

It is intuitive that integrating variation graphs with GWAS should allow for a more thorough exploration of the genotype-phenotype map, and thus support the discovery of associations between DNA sequence variants and traits of interest that are not detectable with SNP-centric analyses. Detailed genotype information coupled with the capacity to represent a wide range of genomic features in variation graphs therefore promises to offer insights that were previously not easily accessible, such as understanding the functional consequences of multi-allelic and complex structural variants.

Conventional GWAS approaches typically rely on genetic variants inferred with the help of linear reference genomes to identify trait associations. This approach has several limitations, particularly when there is a high level of sequence differences between the reference genome and the samples being analyzed. In particular, this can lead to reference bias and the failure to identify causal genetic variations. A powerful alternative is being offered by genome graphs. So far, these were often constructed by anchoring them to a linear reference genome and incorporating known variants identified from reference-based methods, following a paradigm established already 15 years ago (Schneeberger et al. 2009). For example, Liu and colleagues (2020) derived a genome graph for soybeans (*Glycine max*) by detecting structural variants with the MUMmer toolkit using a single reference, and then genotyping almost 3,000 soybean accessions with the vg toolkit. The efficacy of this approach was demonstrated through the identification of a specific structural variant on chromosome 15 that had not been previously linked to seed luster variation by GWAS or other means. Similarly, He and colleagues (2023) developed a pan-genome graph for *Setaria italica* by aligning 112 genomes to a reference and identifying variants with Syri (Goel et al. 2019; Q. He et al. 2023). Nearly 2,000 *S. italica* accessions were then genotyped with this graph and the variants were used in GWAS on 68 traits. Zhou et al. (2022) used a most complex two-step approach to create a tomato genome graph. Initially, they constructed a structural variation graph using HiFi long reads aligned against a reference genome. This graph was expanded by integrating additional variants identified from 706 tomato accessions using DeepVariant for calling indels and SNPs (Poplin et al. 2018), and Paragraph for SV genotyping (Chen et al. 2019). Unlike the other two studies, this method integrated all variants into a unified genome graph, capturing a broader spectrum of genetic diversity and missing heritability (Zhou et al. 2022). Impressive as they were, these studies still had inherent limitations such as relying on reference-based approaches and the potential introduction of bias through variant representation in the VCF format. Additionally, the accuracy of genotyping using short-read alignment often falls short of the precision required for comprehensive variant detection (Du, He, and Jiao 2024).

As alternatives to graph-based approaches, several methods have emerged that completely eliminate the need for linear references. DBGWAS (Jaillard et al. 2018, 2017) is one such method, using an alignment-free, k-mer based approach with compact de Bruijn graphs (cDBG). These represent overlaps between k-mers, removing redundancy, and capturing genetic variations in nodes and edges. Another approach, k-mer GWAS (C. He et al. 2021; Voichek and Weigel 2020; Gaurav et al. 2022), determines the presence or absence of k-mers without relying on reference genomes.To avoid excessive computation for association of all k-mers, which are much more numerous that SNPs, Voichek and Weigel (2020) implemented a prefiltering step that greatly reduces the number of required association tests. While elegant, a major disadvantage of k-mer-based methods is that they are very sensitive to sequencing errors, requiring careful filtering. Additionally, it is difficult to interpret the results without broader genomic context for the detected associations.

To facilitate the incorporation of variation graphs into existing GWAS workflows, we created the tool *gfa2bin*, which converts genome graphs from GFA (Graphical Fragment Assembly) format into a graph-based genotype matrix suitable for GWAS analyses. For seamless integration with existing bioinformatics pipelines, we offer outputs in two different formats: the widely used PLINK format and the more versatile BIMBAM format (Purcell et al. 2007; Servin and Stephens 2005), and we provide a complete Snakemake (Mölder et al. 2021) pipeline of the GWAS alignment-based workflow in the Github repository of *gfa2bin*. Using *Arabidopsis thaliana* as an example, we show that GWAS with variation graphs compares favorably to traditional reference-based approaches, although the best results are obtained when combining multiple methods. It offers unique benefits such as mitigation of reference bias and providing genomic context on the graph structure.

## Results

### General functionality

To enable GWAS with genome graph information, we first create genotype information based on coverage information from sequence-to-graph alignments or from the presence-absence information at multiple levels (e.g., nodes) within the graph. Our newly developed *gfa2bin* tool supports the use of graph alignment coverage from GAM, BAM, or CRAM files to compute a graph-based genotype matrix, and then rapidly and efficiently converts the matrix into the BED (PLINK) format (Purcell et al. 2007), a binary format widely used in GWAS pipelines. The tool also supports output in the BIMBAM format (Guan and Stephens 2008), suitable for imputed genotypes and capable of representing copy-number variation of nodes.

To derive graph alignment coverage information, we implemented techniques for processing sequence-to-graph alignments. In our *A. thaliana* use case, the graph works as a variation-aware reference, taking advantage of the improved mapping qualities compared to traditional references. Essentially, these alignments are converted into coverage information for each base within the graph. *Gfa2bin* uses either plain text or compressed pack files to generate a presence-absence matrix based on a sample-specific dynamic threshold. Nodes equal or above the threshold correspond to reference (REF) or alternative (ALT) alleles in the PLINK format. We arbitrarily coded alternative alleles as 1, and reference alleles as 0. Combined with a kinship matrix - derived from the same dataset or external data - and phenotypes, this is then used as input for any of the available GWAS workflows. Nodes with significant levels of association with specific phenotypes can then be translated into a reference-based representation. This includes both nodes that are directly present in the reference genome, as well as nodes that are not but that are linked to the reference via their closest neighbor node that is present in the reference and that is identified with the *gfa2bin* nearest command. The use of multiple references acts as a convenient sanity check (Figure S3).

Finally, we provide a reference workflow with default parameters to enable rapid adaptation of graph-based GWAS by practitioners in the field (Figure S1).

### GWAS validation

In our *A. thaliana* case study, we first aligned 1,008 short-read datasets (1001 Genomes Consortium 2016) to a genome variation graph containing 28 genomes (Igolkina et al. 2024). We included 1,695 traits with multiple testing correction using hundredfold permutation for each trait. With the graph nodes, we found significant associations for 637 traits, compared to 668 traits with k-mers, and 594 traits with SNPs. The largest class comprised the 398 traits that had significant associations with all three inputs (Figure 2A). The largest overlap between exactly two inputs was observed for k-mers and graph nodes, 129, while SNPs had the highest number of unique hits, 88.

**Figure 1.**
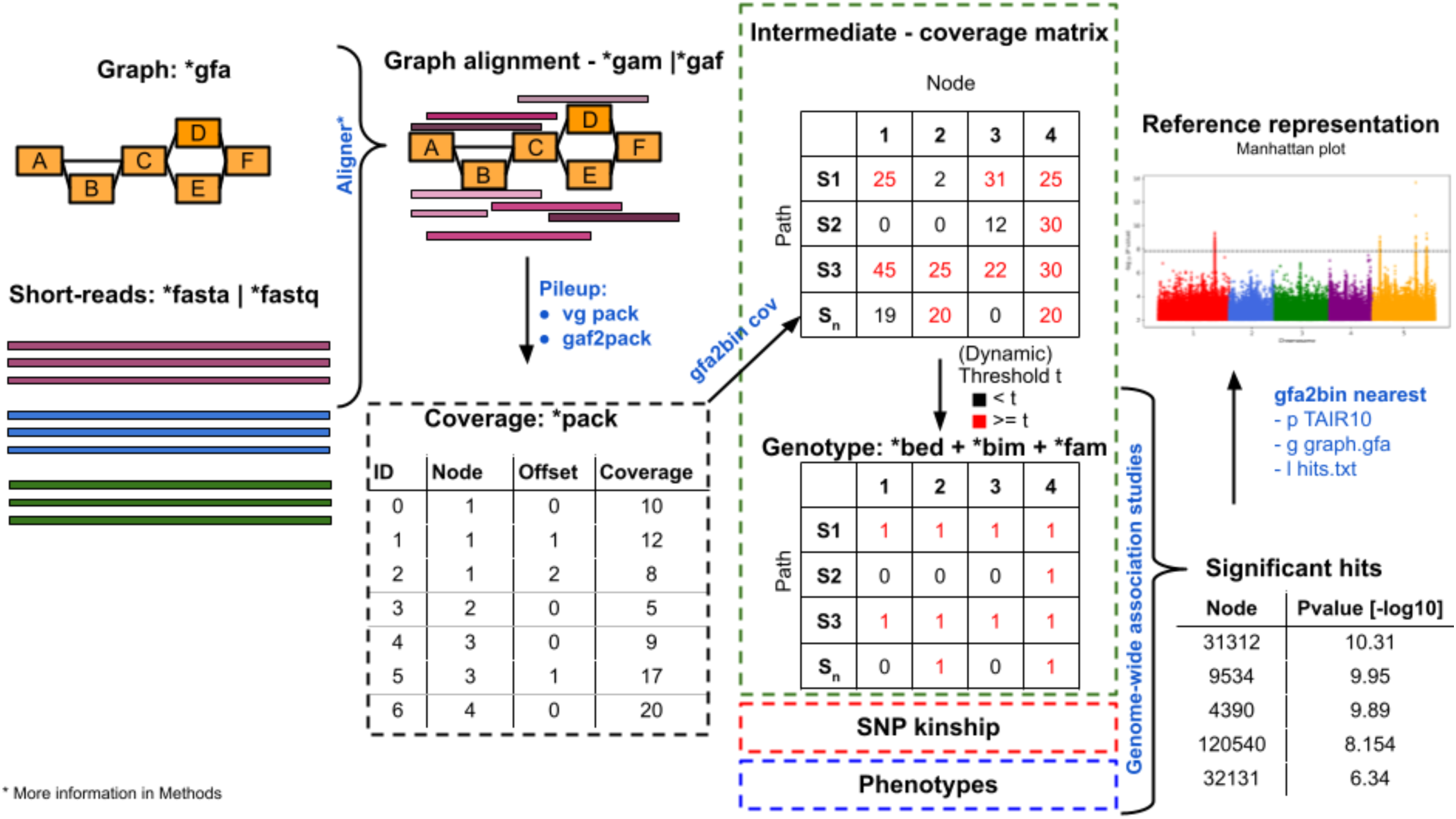
Schema of the graph-alignment GWAS workflow. Read data from multiple samples are aligned to a graph in gfa format, followed by calculation of the coverage per node in each sample. Subsequently, the coverage matrix is converted into a binary genotype matrix using dynamic thresholding. Thos genotype matrix, a kinship matrix and phenotype data are provided as inputs to a GWAS workflow, such as *GEMMA*. Nodes with significant GWAS results can be mapped back to a linear reference representation, which can then be shown as a Manhattan plot, or directly visualized on a subgraph (not shown here, see Figure 3B).

**Figure 2.**
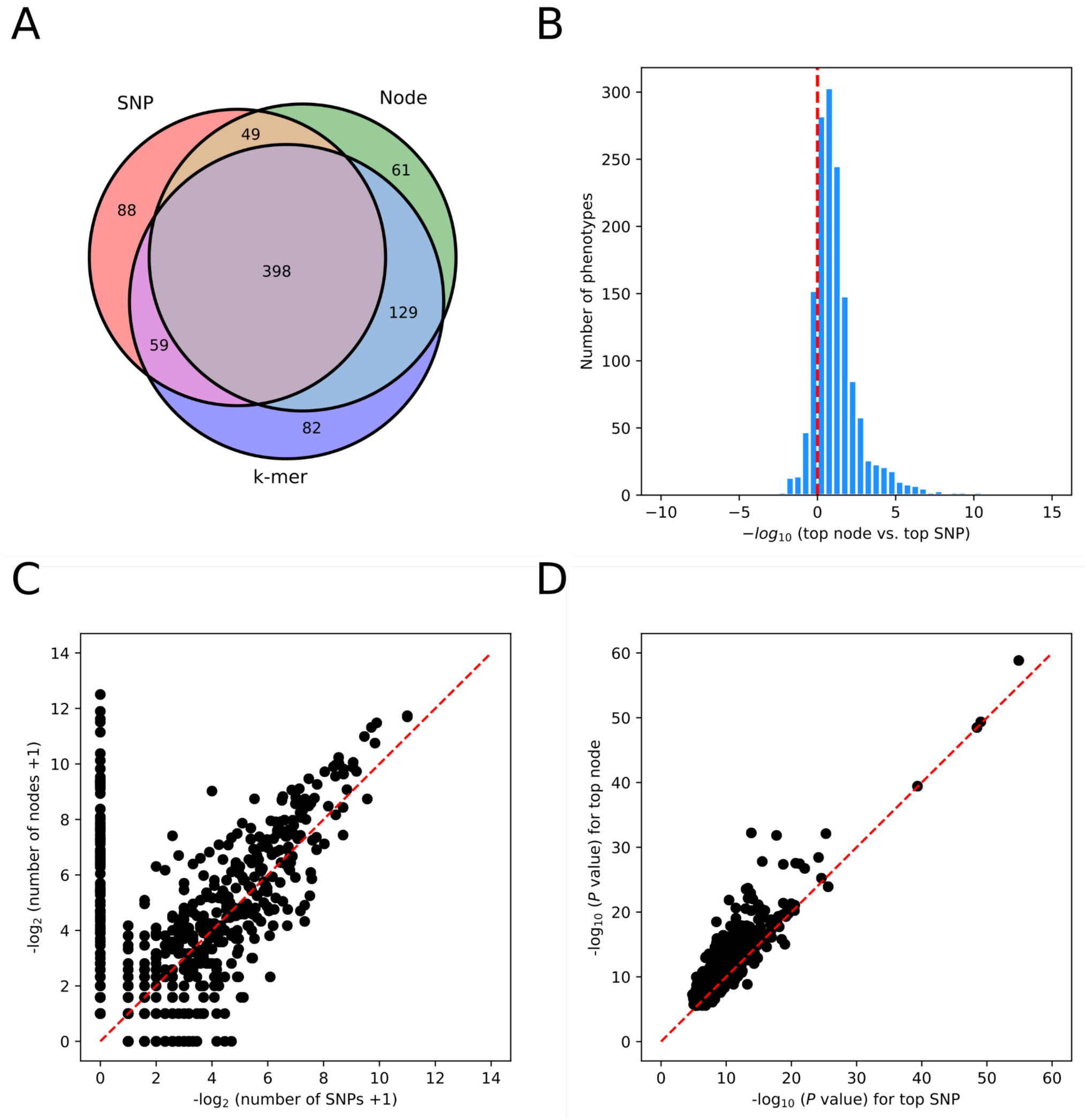
Comparison of SNP-, k-mer-, and graph node-based GWAS on 1,695 *A. thaliana* phenotypes. (A) Overlap between phenotypes with SNP, k-mer and graph node hits. (B) Ratio between top P values (expressed as -log10) for graph node and SNP hits. (C) Correlation of numbers of significant graph nodes versus SNPs. (D) Correlation of P values of top nodes with SNPs (r=0.91).

In a direct comparison to SNPs, the use of graph nodes as input identified a greater number of significant hits in shared associations, along with a higher incidence of hits unique to the graph methodology (Figure 2C). Additionally, these graph node-based hits had on average lower P values, particularly when focusing on the top hit for each trait (Figures 2B and 2D). The high correlation and the positive long-tailed distribution of the ratios between P values from graph nodes and SNPs (see Figures 2B and 2D) validated our approach and the robustness of the underlying graph mapping for GWAS. In traits common to both inputs, we did not detect any exceptional outliers.

For a more in-depth validation, we used flowering time, which is a thoroughly investigated trait in *A. thaliana* (Cho, Yoon, and An 2017). Our graph node approach successfully re-identified several known associations (1001 Genomes Consortium 2016), including with the loci *FLOWERING LOCUS T* (*FT*), *FLOWERING LOCUS C* (*FLC*), *DELAY OF GERMINATION 1* (*DOG1*) and *VERNALIZATION INSENSITIVE 3* (*VIN3*) (Figure 3A). Direct visualization and re-annotation revealed an insertion in *FLC*, depicted as a loop in the graph representation of genome sequences, as being responsible for the detected association with flowering time in our dataset (Figure 3B). The linear representation of genome sequences in Figure 3C confirmed the indel nature of the polymorphism, which is absent from the majority of our 28 reference genomes, including the original TAIR10 reference genome. The paths traversing the loop, which correspond to four of our 28 reference genomes in Figure 3C, are associated with later flowering (Table S1).

**Figure 3.**
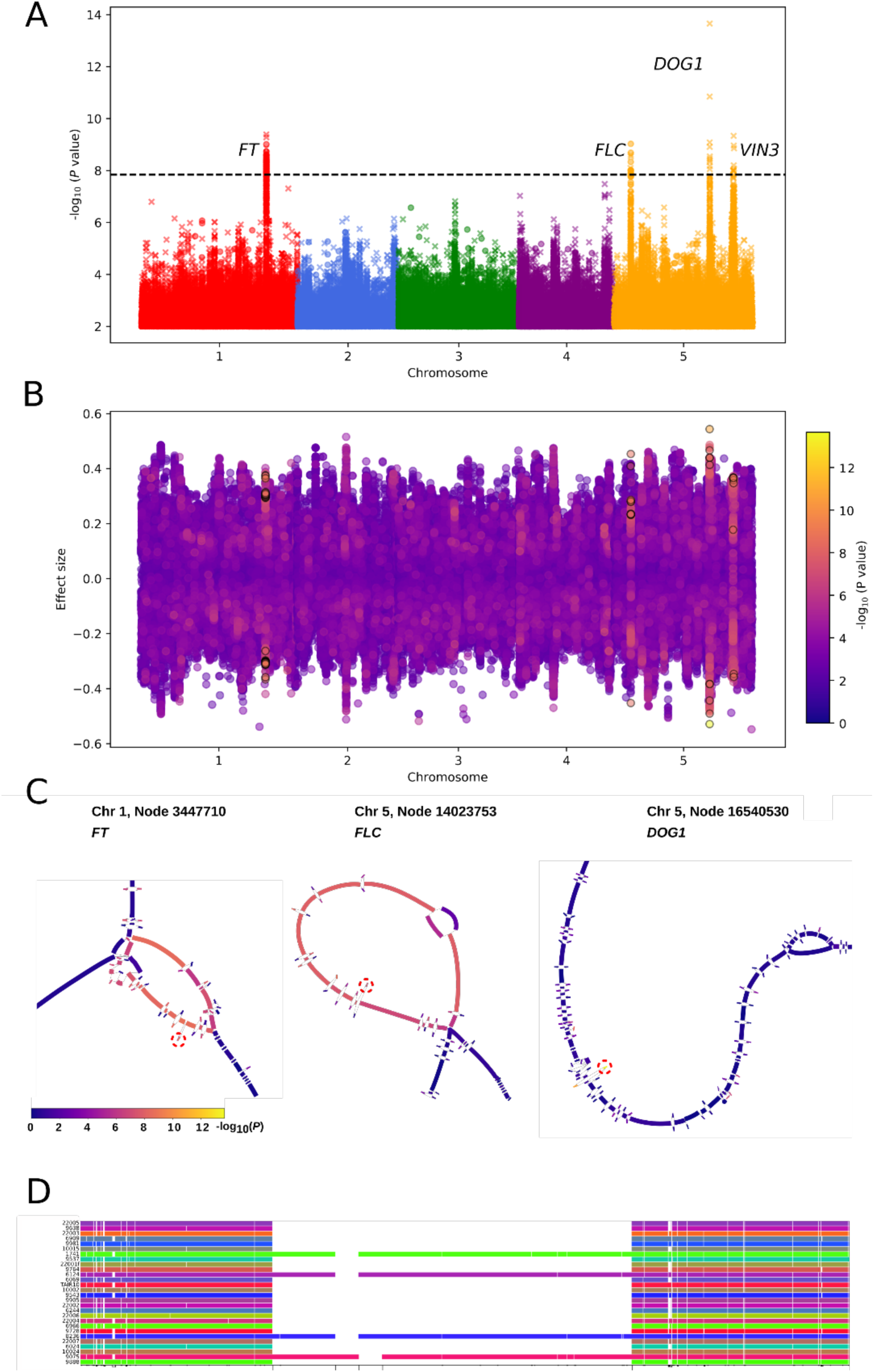
Interpretation of node-based GWAS. (A) Associations of graph nodes with flowering time at 10°C (FT10) as trait, shown on a linear representation after conversion to the TAIR10 standard reference. Nodes present in the reference are denoted by **x**. Nodes that are absent from the reference are denoted by **o**. They were linked to the reference by finding the closest reference node, as described in the Methods. (B) Effect size of each node (linked to TAIR10 position) computed by point-biserial correlation and without kinship correction. Significant hits are highlighted with black outlines. (C) Subgraph visualization for associations with FT (4912 bp shown), FLC (2355 bp shown) and DOG1 (1336 bp shown). Note the loop structure at FLC, which identifies sequences absent from the TAIR10 reference. (D) Representation of the FLC subgraph on a linear (graph node) scale.

We calculated the effect size of presence/absence of nodes on flowering time at 10°C (FT10) without kinship correction. While the significant GWAS hits did not necessarily have the largest effect sizes – indicating the importance of kinship correction (Figure 3B), it was noteworthy that significant nodes at a locus included ones with opposite effects, indicating that our approach identifies what appear to be “alternative” alleles.

A similar bubble-like structure was also observed for the *FT* locus (Figure 3C). Even though the nodes along the whole structure were associated with elevated P values, the most significant graph node was found in one of the alleles with a size of 1 bp, indicating a short indel polymorphism.

The *DOG1* locus showed a linear structure, with the most significant node representing a single SNP, comparable to traditional SNP-based GWAS (Figure 3D).

### Additional workflows

We developed a comprehensive Snakemake pipeline that guides users through all necessary steps for graph node GWAS, from raw sequence data to generation of GWAS inputs in PLINK format. This end-to-end solution ensures a smooth and standardized process, reducing potential errors and inconsistencies in data preparation. The requisite coverage vector can be derived from almost all graph alignment formats, providing users with the flexibility to use their preferred mapping software.

Although our alignment workflow also works with plain-text coverage data, we provide a custom-built packing tool, designed to reduce disk-space and facilitate efficient preprocessing. The preprocessing step incorporates normalization techniques, if desired.

### Other functions

To enhance user experience and expand the functionality of *gfa2bin*, we incorporated several pre- and post-processing features. Notably, *gfa2bin* includes a ’view’ function that allows users to convert their PLINK files into VCF format, thereby increasing compatibility with a range of genomic analysis tools. Additionally, users can directly remove or filter sample genotypes from PLINK files using the ’remove’ and ’filter’ functions. For post-analysis interpretation, we developed the ’find’ and ‘nearest’ function within *gfa2bin*. This feature processes a list of significant or important nodes, such as those identified through GWAS, and returns their exact locations within each genomic path, or in the case of ‘nearest’, the closest reference position. This functionality is crucial for contextualizing results, enabling researchers to swiftly locate and investigate regions of interest within the complex genome graph.

To ensure broad compatibility with different GWAS tools, our PLINK BED format output includes a diploid representation for each sample. This feature is essential for accurately modeling genetic variation across different organisms, and supports analyses that account for heterozygosity and allele-specific effects.

## Discussion

We have shown the *gfa2bin* tool to be an effective and versatile solution for converting variation graphs to GWAS-ready formats. Starting from alignment of short reads to a graph that captures most common structural variants in the global *A. thaliana* population, we reproduced many associations that were identified before using conventional GWAS approaches with a single linear reference genome (1001 Genomes Consortium 2016). Going beyond SNP-based GWAS, graph-based GWAS identified more associations with greater statistical significance, but there were also associations that were identified only with SNPs or only with k-mers. This indicates that the improved representation of genetic diversity through variation graphs enhances the scope and effectiveness of GWAS, but that the best results are obtained by combining multiple methods. Our workflow allows for straightforward interpretation of graph node hits in the context of a standard reference genome, which in turn facilitate integration of results with those from SNP-based approaches. A drawback of k-mers is that k-mers not present in the standard reference genome can be difficult to interpret (Voichek and Weigel 2020). In principle, it should be possible to link k-mer hits to graph node-based hits by mapping k-mers to the genome graph.

Associations found only with SNPs can at least in some cases be traced back to graph nodes containing long sequences (Figure S2). We interpret this as the causal SNP not being captured in our set of 28 genomes used to construct the graph. If a single SNP is found in a longer node, this would only marginally reduce coverage of the node by the sampled short reads, and we would score the node as present regardless of the SNP. Conversely, multiple alternative alleles nor represented at a node in the graph, might all be scored as zero coverage. This would reduce power in association testing, if only some of these alternative alleles lead to phenotypic differences.

Our tool supports genotyping at base pair level, which, while computationally intensive, would aid in uncovering more associations and mitigating the effects of averaging factors. In addition, a larger number of diverse genomes to build the initial graph for short read mapping should provide for a greater set of nodes that can be tested for association with different phenotypes.

We not only provided a framework for coverage and node-based GWAS in variation graphs, but also support the usage of complex *pggb* graphs, which have not yet been used for mapping-based GWAS approaches. These graphs are designed to incorporate alignments of all input sequences and do not rely on pre-computed variation.

In the long term, to fully exploit all of genomic complexity, a more sophisticated representation of variants in graphs might be necessary. This would include the ability to represent and identify both simple and complex structures, or combining multiple nodes into, e.g., bubbles with different alleles. Additionally, at the alignment level, a more nuanced representation of potential alternative alleles (variants), which may or may not be artifacts of the alignments, could enhance the identification of associations.

It is crucial to recognize that, as in traditional SNP-based GWAS workflows, input quality is a primary determinant of ability to detect associations. As with SNP-based workflows, coverage depths and error rates in the short read data set used to call variants on the graph are one such determinant. In addition, the quality of the variation graph used for mapping will significantly influence the GWAS results. On the one hand, quality is determined by the number and accuracy of complete input genomes for building the graph. On the other hand, choices made when building the graph must be carefully considered, because the optimal graph will depend on the extent of variation present in the input genomes (Garrison et al. 2023).

In future, instead of mapping short reads to a graph, one could use completely assembled genomes to genotype the paths in the graph. This would not only obviate the need for arbitrary thresholds to call node presence and absence, but would also allow for the use of copy number variants, inferred from the number of times a specific genome path visits the same node, in GWAS. In anticipation of this bright future, we have already implemented commands that could accommodate such analyses.

## Materials and methods

### Datasets

The datasets were sourced from published resources. *Arabidopsis thaliana* assemblies for graph construction were obtained from the 1001 Genomes Plus project (Igolkina et al. 2024). Short reads were from the 1001 Genomes Project (1001 Genomes Consortium 2016) and corresponding phenotypes from multiple publications, collected and curated in (Voichek and Weigel 2020).

### Kinship matrix

To facilitate the comparison between SNP-, k-mer and graph node-based GWAS, we used the same kinship matrix for all three. The kinship matrix was originally calculated on SNP data using the method from EMMA (Kang et al. 2008; Voichek and Weigel 2020).

### Genome graph building

The *A. thaliana* graph was constructed with *pggb* (Garrison et al. 2023). The *pggb* workflow was executed with *wfmash* (v0.10.2-2-gb310bd1), *seqwish* (v0.7.8-3-gd9e7ab5), *odgi* (v0.8.2-92-gbfae0b3), and *smoothxg* (v0.6.8-31-g06bbf35). Our parameter set was -s 10000 -k 79 -n 27 -p 90 -P asm10.

### Graph mapping of short reads

Short reads were mapped to the graph with a custom pipeline (Figure S1), provided in our Github repository. The reference sequences are extracted directly from the graph, then short-reads were mapped to the resulting “super-genome” using *bwa mem* and converted to BAM format by *samtools*. We transferred the BAM files to Graph Alignment / Map (GAM) format using the mapping relationship between the input sequences and the genome graph. Coverage information was calculated using *vg pack.* Finally pack coverage was converted to pack compressed format to reduce storage using the newly implemented packing tool.

Alternatively, users can convert the linear alignment to the Graph-aligned format (GAF) and convert to coverage information using our custom tool *gaf2pack* (https://github.com/MoinSebi/gaf2pack).

Without the “super-genome”, sequences can be aligned directly to the graph using established mappers such as *vg giraffe* (Sirén et al. 2020) or *graphaligner* (Rautiainen and Marschall 2020). The resulting alignment can be directly converted to pack format. The provided pipeline supports pack files as input to the GWAS workflow (Figure S1).

### Converting nodes to reference positions

We anchored all nodes tested in GWAS using *gfa2bin nearest* to specific locations on the TAIR10 reference genome. Specifically, we identified the shortest path (in base pairs) to any reference node and recorded the distance in base pairs. To denote the proximity to the respective anchor node in the reference, different markers or colors can be used in Manhattan plots (Figure S3).

### Visualization

Manhattan plots were generated using the *Python matplotlib* library. To enhance clarity, we excluded all hits falling above a threshold of log_10_(−2). Graphs were represented in two dimensions (“2D”) using Bandage NG (Wick et al. 2015; Korobeynikov, n.d.). The graph layouts show the significant node and its 50 neighboring nodes. For a linear (“1D”) representation of the *FLC* region in the graph, *odgi viz* was employed. We extracted a subgraph using *Bandage NG* and manually added the represented path to facilitate detailed visualization.

### Effect size

For effect size computation we used Point-Biserial Correlation. We selected this correlation technique to highlight extent and direction (positive or negative) of the relationship and get insights into whether two groups differ in terms of their average outcome.

### GWAS validation

To compare the results from different GWAS inputs, we followed a published workflow (Voichek and Weigel 2020). This includes ranking nodes by their initial scoring, then using the 10,000 top ranked genotypes to calculate the exact P values, for which a permutation-based threshold is calculated. This process is repeated until all hits are in the top half of the initial ranking, passing more entries with each repetition.

SNP and k-mer hits for the set of phenotypes examined have been published (Voichek and Weigel 2020). Among the shared associations, we filtered out traits with fewer than 40 phenotyped accessions, or where there was a difference between SNP GWAS and a uniform distribution (Kolmogorow-Smirnow test, p ≤ 0.05; adapted from (Voichek and Weigel 2020)).

## Acknowledgements

We thank Christian Kubica and Erik Garrison for helpful discussions.

## Funding

This work was supported by the Novo Nordisk Foundation (Novozymes Prize) and the Max Planck Society.

## Author contributions

S.V., I.B., Z.B., W.X and D.W. conceptualized the study and conceived the tool functionality and analyses; S.V. wrote the gfa2bin code; I.B. wrote the Snakemake pipeline; S.V. and I.B. performed the analyses; S.V., I.B. and D.W. wrote the manuscript with input from all authors.

## Data and code availability

Source code is available under MIT license at https://github.com/MoinSebi/gfa2bin.

## Competing interests

D.W. holds equity in Computomics, which advises plant breeders. D.W. also consults for KWS SE, a plant breeder and seed producer with activities throughout the world. The other authors declare no competing interests.

## Notes

https://github.com/MoinSebi/gfa2bin

