## Supplementary Information for "Gfa2bin enables graph-based GWAS by converting genome graphs to pan-genomic genotypes"

### 1. Implementation

**G*fa2bin*** was implemented in the  programming language Rust, incorporating several Rust crates to enhance performance and enable multithreading. An additional tool, ***packing***, for reducing storage of coverage files was developed. ***Packing*** can be used for compression and/or normalisation of coverage information.

#### A. Inputs

The alignment-based workflow utilises coverage information as input, which can be provided in either plain-text pack format, or in compressed or normalised formats generated by the packing method. Pack files are typically produced by either **gaf2pack** (https://github.com/MoinSebi/gaf2pack) or **vg pack**, which is part of the variation graph framework (Garrison et al. 2018). Depending on the alignment algorithm, the inputs are either in Graph Alignment Format (GAF) or Graph Alignment Map (GAM) format. Further details on graph alignment are provided below.

When using our graph subcommand, only a graph in GFA format is required. We do not support any conversion for rGFA. Other methods of our tools rely on a genotype table (PLINK bed, bim, and fam files) in conjunction with the associated graph structure. Additional details on alternative methods are listed below (under 'Additional Methods').

#### B. Thresholds

We provide multiple options for computing thresholds for each sample in our tool. In addition to a fixed threshold applied across all samples, we recommend using dynamic calculations to account for varying coverage levels across samples. Metrics that do not rely on distribution assumptions, such as the median or percentiles, generally provide a more accurate representation than the mean. For each metric, we offer the ability to apply a scaling factor, which multiplies the calculated value by a user-defined number. An example is provided below.

#### C. Outputs

The primary output of our method is the PLINK BED format, which supports diploid binary representation of multiple samples and genotypes. Diploid features are available only in our *graph* methods, specifically through **panSN-spec** (https://github.com/pangenome/PanSN-spec). Our alignment method, however, does not support diploid representation, as each dataset corresponds to a single sample. Furthermore, the coverage information does not include detailed processing of homozygous or heterozygous samples, which may limit its applicability in high heterozygosity diploid organisms.

For certain analyses, we also provide output in BIMBAM format. While BIMBAM was originally developed for imputed data, we use it here to represent copy number variation (CNV) of nodes, such as when a single sample contains multiple copies of the same nodes, or in cases of 'over-coverage' of single nodes during alignment-based runs.

#### D. Methods overview

This list provides a brief summary of all methods available in our tool. More detailed information can be found in our GitHub repository.

##### Creating Genotypes:

- **Cov**: Converts coverage information into genotypes, using the graph as a variation-aware reference during alignment.
- **Graph**: Generates genotypes based on the presence or absence of nodes, directed nodes, or edges for each path in the graph.
- **Subpath**: Starting from a specific node, this method collects all subpaths for a given maximum step number that traverse the node based on path information. The presence or absence of each path within each subpath group is compared to construct genotypes for each node in the graph.
- **Window**: Builds new genotypes from the binary genotype matrix by including a window of neighbouring N genotypes for each existing genotype. The window size is adjustable, as it plays a crucial role in genotype formation.

##### Modification:

- **View**: Converts PLINK (bed, bim, fam) files into VCF format.
- **Merge**: Merges multiple PLINK files, assuming they contain the same samples in the same order. This is often used to combine a previously split dataset.
- **Split**: Splits a PLINK file into multiple files of equal size.

##### Post-processing:

- **Find**: Locates the exact position of a specific node within any path in the graph, returning the start and end intervals of the node.
- **Nearest**: Identifies the nearest (or closest) node on the reference path.

**Table S1. Geographic and flowering time of accessions used for graph construction.**

This table presents the geographic coordinates (longitude and latitude) and flowering time data for the *A. thaliana* accessions used to build the graph (Igolkina et al. 2024). Orange highlights accessions with the *FLC* insertion shown in Figure 3 of the main text.

| Accession ID | Flowering time | Longitude | Latitude | Country |
|---|---|---|---|---|
| **1741** | - | -85.398 | 42.405 | US |
| **6024** | 106.75 | 13.37 | 55.75 | Sweden |
| **6069** | 107.25 | 18.28 | 62.95 | Sweden |
| **6124** | 129.50 | 13.31 | 55.84 | Sweden |
| **6244** | 90.75 | 18.47 | 62.92 | Sweden |
| **6909** | 70.50 | -92.30 | 38.30 | US |
| **6966** | 62.25 | -0.64 | 51.41 | United Kingdom |
| **8236** | 74.00 | 15.76 | 49.33 | Czech Republic |
| **9075** | 93.25 | 48.61 | 38.74 | Azerbaijan |
| **9537** | 64.25 | -6.66 | 38.07 | Spain |
| **9543** | 94.33 | -5.39 | 36.77 | Spain |
| **9638** | 88.25 | 80.86 | 51.73 | Russia |
| **9728** | 77.50 | 18.90 | 48.46 | Slovakia |
| **9764** | 69.75 | 35.84 | 34.10 | Lebanon |
| **9888** | 86.25 | -3.31 | 40.93 | Spain |
| **9905** | 89.75 | -4.01 | 40.76 | Spain |
| **9981** | 69.25 | 16.24 | 38.76 | Italy |
| **10002** | 61.00 | 9.04 | 48.53 | Germany |
| **10015** | 69.75 | 71.30 | 37.29 | Afghanistan |
| **10024** | - | 36.21 | -2.87 | Tanzania |
| **22001** | - | 115.06 | 32.14 | China |
| **22002** | - | 108.61 | 27.94 | China |
| **22003** | - | -4.10 | 34.09 | Morocco |
| **22004** | - | -7.41 | 31.47 | Morocco |
| **22005** | - | -17.13 | 32.75 | Portugal |
| **22006** | - | -16.93 | 32.74 | Portugal |
| **22007** | - | 38.06 | 13.24 | Ethiopia |

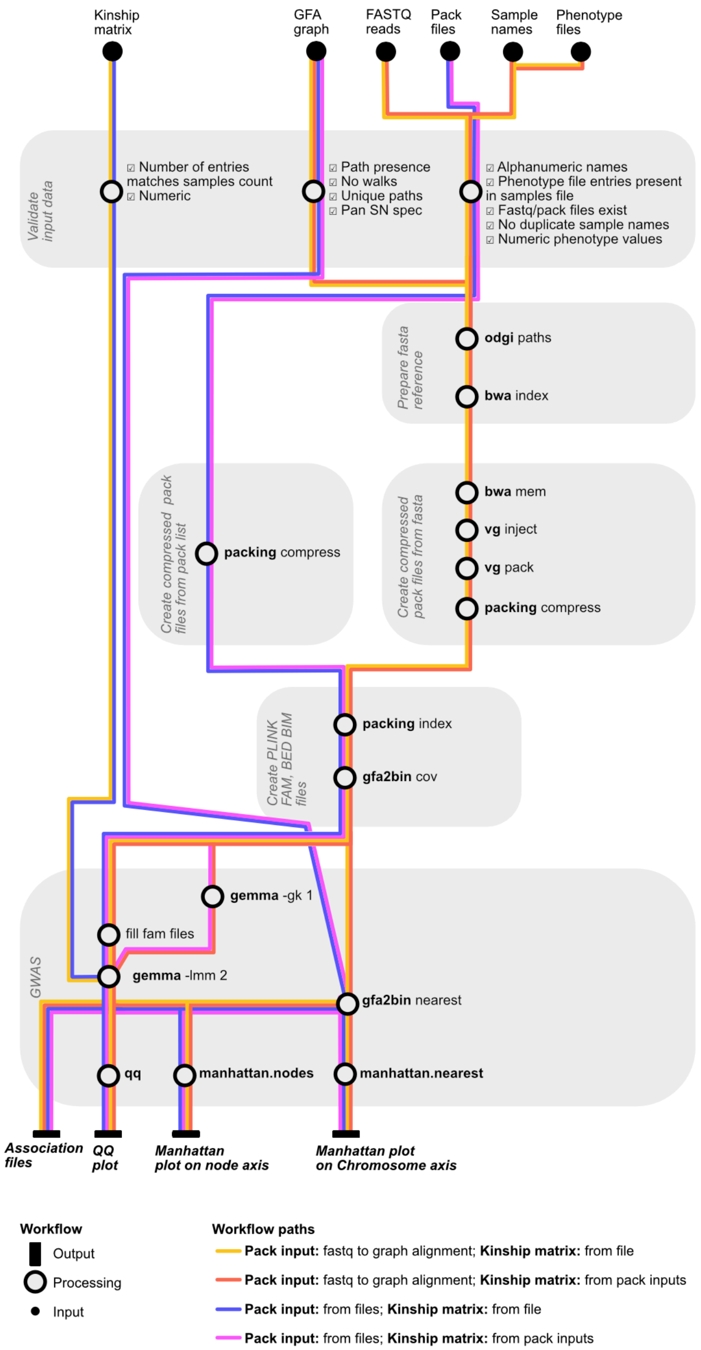


**Figure S1. Diagram of the graph-based workflow as implemented in the provided pipeline.**

Sequences can be aligned linearly to all genomes simultaneously, with linear alignments translated into graph terms using the graph structure itself. This conversion facilitates the calculation of coverage by the graph alignment. Alternatively, externally generated pack files can be processed.

A

B

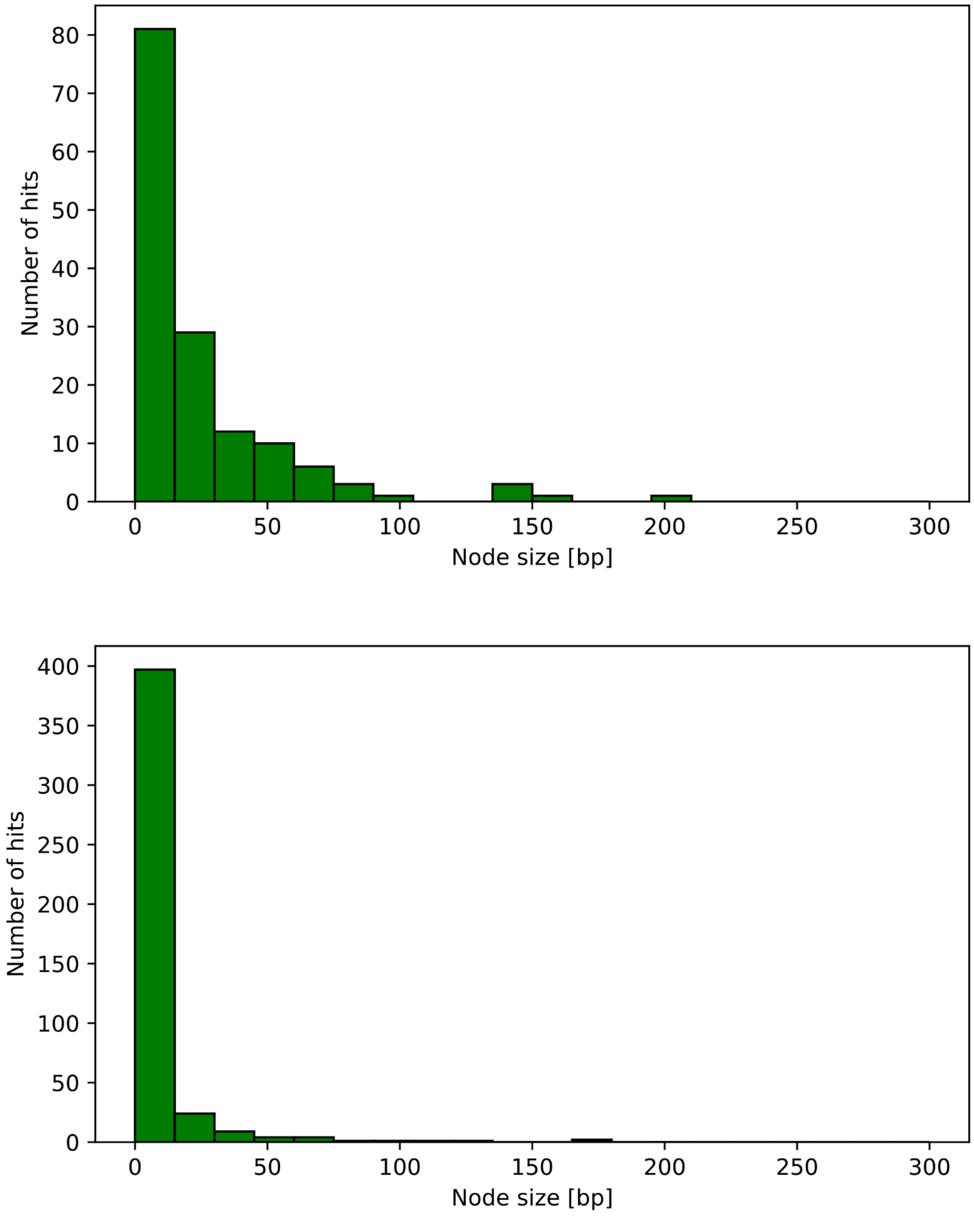


**Figure S2. Node sizes of the top hits.**

(A) SNP-exclusive GWAS hits, transferred to the graph-coordinate system (nodes). (B) Shared SNP- and graph node-based hits. SNP-specific hits are mostly found in longer nodes with a median of 11 bp compared to 1 bp in the shared dataset.

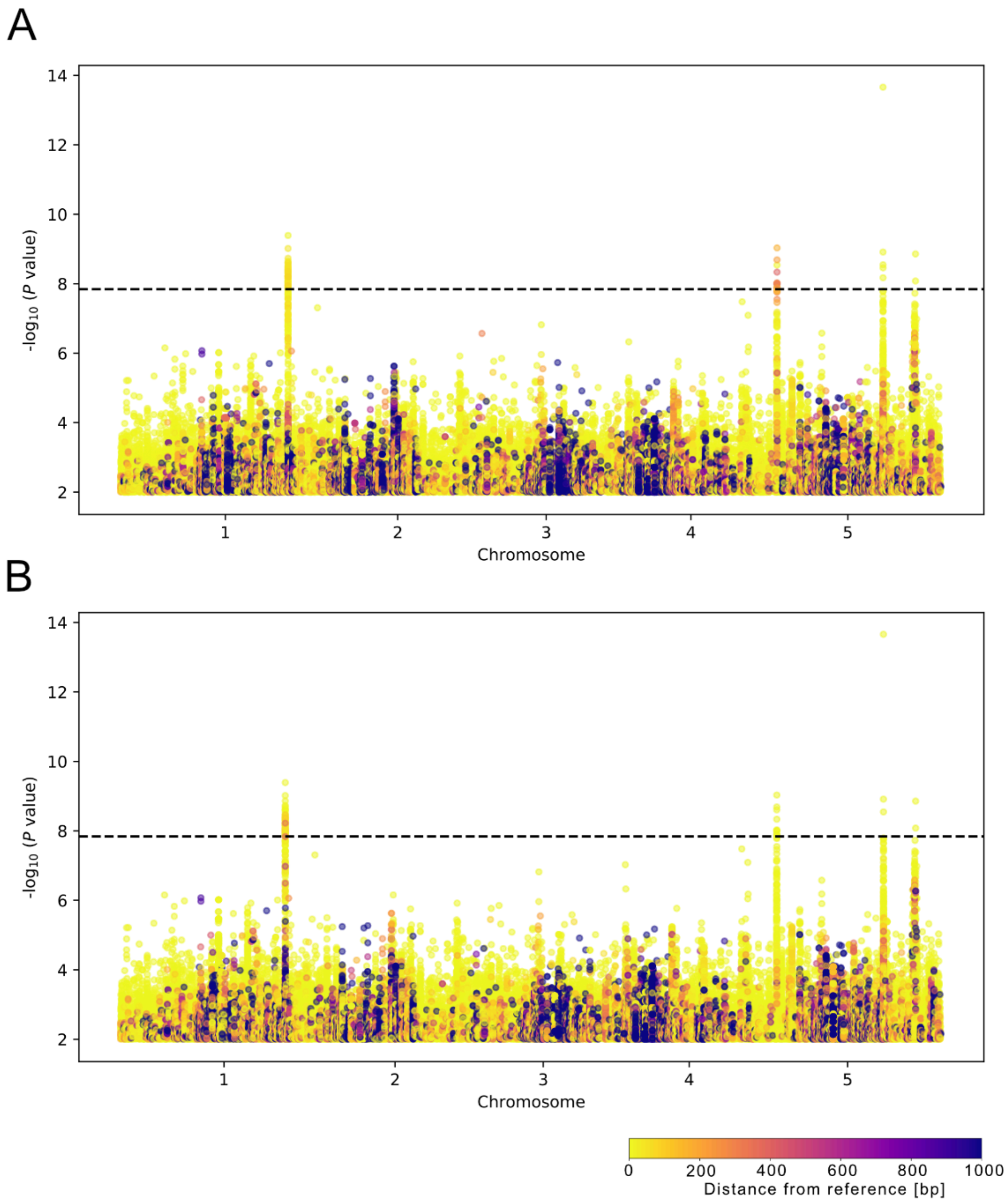


**Figure S3. Manhattan plots for FT10 trait projected onto different reference genomes.**

Graph nodes were mapped to (A) TAIR10 or (B) KBS-Mac-74 (accession ID 1741). Overall results are similar, underscoring the robustness of the approach.
